# Linking Structure and Function: Image-Based Virtual Populations of the Retinal Vasculature

**DOI:** 10.1101/2023.12.05.570054

**Authors:** Rémi Hernandez, Savita Madhusudhan, Yalin Zheng, Wahbi K. El-Bouri

## Abstract

**Purpose:** This study explores the relationship between microvascular parameters as delineated by optical coherence tomography angiography (OCTA) and retinal perfusion. We introduce a versatile framework to examine the interplay between the retinal vasculature structure and function, generating virtual vasculatures from central retinal vessels to macular capillaries. Alongside this, we develop a haemodynamics model which tests the associations between vascular morphology and retinal perfusion.

**Methods:** The generation of the vasculature is based on the distribution of four clinical parameters pertaining to the dimension and blood pressure of the central retinal vessels, constructive constrained optimisation and Voronoi diagram. Arterial and venous trees are generated in the temporal retina and connected through three layers of capillaries at different depths in the macula. The correlations between total retinal blood flow and macular flow fraction and vascular morphology are derived as Spearman rank coefficients and uncertainty from input parameters is quantified.

**Results:** A virtual cohort of 200 healthy vasculatures were generated. Mean and standard deviation for retinal blood flow and macular flow ratio were 19.15 *±* 7.34 μL*/*min and 4.52 *±* 1.19 %. Retinal blood flow was correlated with vessel area density, vessel diameter index, fractal dimension and vessel calibre index. The macular flow fraction was not correlated with any morphological metrics.

**Conclusions:** The proposed framework is able to reproduce vascular networks in the macula that are morphologically and functionally similar to real vasculature. The framework provides quantitative insights into how macular perfusion can be affected by changes in vascular morphology delineated on OCTA.

## 1 Introduction

The retina is a highly oxygen-dependent, complex band of tissue at the back of the eye which plays a key role in visual function. It requires close interplay between many different cell types and supporting structures, including a complex vascular system^32^. As a result, the retina is sensitive to small changes which may lead to loss of visual functions. *In silico* modelling has the potential to offer insight into the complex interactions between the retinal environment and the underlying causes of retinal diseases. In particular virtual populations and *in silico* clinical trials are a promising way to enhance basic research and clinical trials^32^. Optical coherence tomography angiography (OCTA) is a non-invasive, new imaging modality that offers three dimensional, high resolution angiograms of the macula, which is the central-most area of the retina. Several microvascular metrics, such as vessel density and fractal dimension, have been suggested to quantify the quality of the microvasculature on OCTAs^17^. Using those metrics, several microvascular changes have been not only linked with ageing and diseased retinae^22;39;55;56;60^ but also with cerebrovascular changes and cardiovascular diseases^28;38;45^. For the latter two, virtual populations have been developed, but similar work is yet to be done for the retinal vasculature or for the linked cerebral-retinal vasculature^32^.

Such alterations to the retinal and choroidal vasculature are expected to negatively affect the perfusion of the retina. Ischaemia and hypoxia are likely involved in the pathogenesis of several retinal diseases, in particular neovascular diseases such as neovascular age-related macular degeneration, proliferative diabetic retinopathy or retinal vein occlusion which are characterised by pathological growth of blood vessels through angiogenesis^40^. Hypoxia is a known trigger of angiogenesis and targeting ischaemia-induced angiogenic pathways might improve treatment outcomes^57^.

However, owing to the difficulty of experimentally measuring oxygen levels in the retina, the role of hypoxia in the pathogenesis of neovascular diseases remains unclear. Furthermore, it also remains unclear how changes in microvascular metrics computed on OCTAs relate to the quality of blood perfusion. Computational models of the retina and its vasculature, combined with mathematical models of haemodynamics and oxygen transport, can help to link structure and function.

Retinal haemodynamics and oxygenation have received considerable attention from the modelling community in recent years^6;11;14;16;19;24;29;32;53;62;64^. Compartmental models^6;16;29^ or symmetrical branching networks^16;53;64^ are used in many of these models. These approaches are favoured for their simplicity and adaptability to systems with limited information. However, they fail to reproduce the complexity and heterogeneity of the retinal vasculature^62^. In contrast, models based on vascular networks reconstructed from imaging data^11;24^ are more faithful to the physiology of the retina, but the reconstruction of the network is arduous and therefore only a limited number of eyes can be modelled. Space-filling algorithms are a way to circumvent these problems by generating heterogeneous networks with similar characteristics to real vasculature^14;35;54^. For instance, Causin et al.^14^ used diffusion-limited aggregation because it creates structures with a fractal dimension similar to that of retinal vasculature. The class of space-filling algorithms called constrained constructive algorithms (CCO) is another approach that includes rules and constraints meant to reproduce the angiogenesis process^35;54^. It has been applied to the retinal vasculature^12;37^, but only to create synthetic data for deep learning applications.

Vascular morphology has been established as a biomarker for the development, progression and prognosis of several retinal diseases^10;61^, including diabetic retinopathy^26;30^ and agerelated macular degeneration^44;55^. Changes in these metrics may indicate impairment to retinal or macular blood flow, which could contribute to the development of the disease. However, quantifying these impairments in a sufficiently large population is challenging with conventional experimental techniques. The modelling framework presented here can quantify these impairments in large virtual populations. The model captures the complexity of the macular vasculature and is able to link imaging biomarkers with physiological parameters. Our study sought to: 1) develop a method for generating cohesive vascular networks in the retina and macula, adaptable to virtual population generation; 2) propose a mathematical model of blood flow in the virtual vasculatures; 3) quantify associations between macular vascular morphology and haemodynamics parameters in a healthy retinal population. Our method constructs a vascular network spanning the superficial layer of the temporal retina, originating from the central retinal artery (CRA) and terminating at the central retinal vein (CRV). In the macula, all three capillary layers are modelled. Additionally, global sensitivity analysis on the method’s hyperparameters lays the foundation for generating tailored retinal populations based on the distribution of morphological metrics.

In Section 3.1, we show a comparison of the microvascular structure of the model with OCTA measurements and validate the blood flow model against two independent experimental studies. In Section 3.3, global sensitivity analysis techniques are applied to each aspect of the model to quantify the uncertainty brought about by the uncertainty in clinical parameters and model hyperparameters. Section 4 contains a discussion of the results, while Section 5 summarises the conclusions and future perspectives.

## 2 Methods

In this section we describe the proposed models. Section 2.1 describes the model that generates the retinal vasculature, from the macro to micro scale. Section 2.2 describes the proposed haemodynamic model.

### 2.1 Structural model

The structural model generates retinal vasculature from the macro scale (arteries, arterioles, veins and venules) to the micro scale (capillaries).

Macroscale vasculature is generated on the temporal retina, starting from the CRA and ending in the CRV. First, a statistical shape model^18^ of the major temporal arcades was developed using a fundus photographs dataset. The remaining superficial temporal vasculature is partially generated with a constructive constrained optimisation (CCO) algorithm^54^.

Microvasculature is generated in the macula area (see Figure 1) across three vascular layers, namely the superficial vascular plexus (SVP), intermediate(ICP) and deep capillary plexuses (DCP), arranged as parallel, planar layers at fixed depths *z*. In contrast, the macrovasculature is only generated in the SVP. This is because the ICP and DCP are composed of capillaries in the perifovea and merge with the SVP outside the macula^3^.

**Figure 1:**
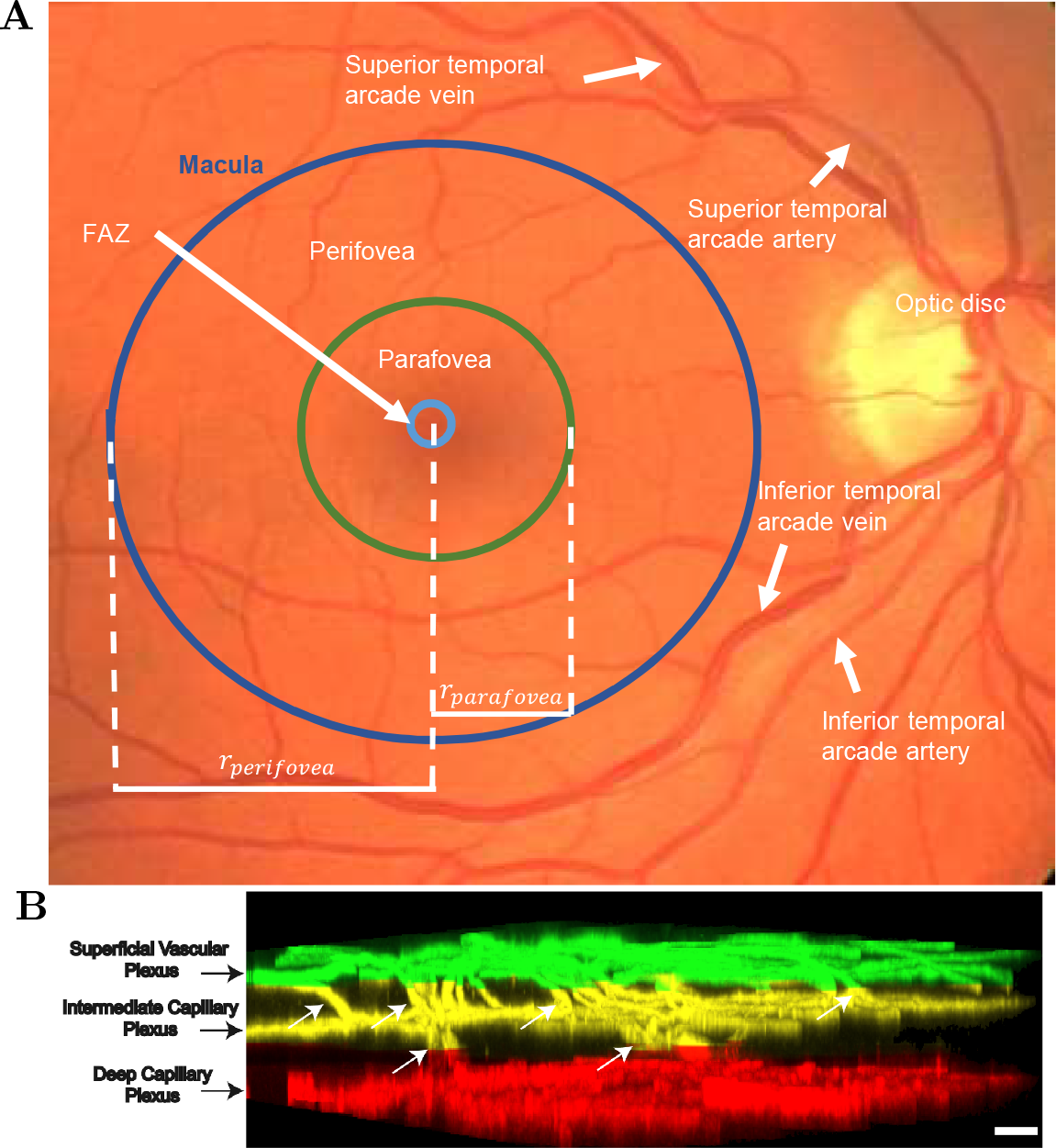
Architecture of the retina. (**A**) Landmarks of the retina on a 45^*°*^ field-of-view colour fundus photograph from the DRIVE dataset^52^. (**B**) The three capillary layers of the macula imaged with histology. Image from the work of^4^, available under a CC BY-NC-ND license.

**Figure 2:**
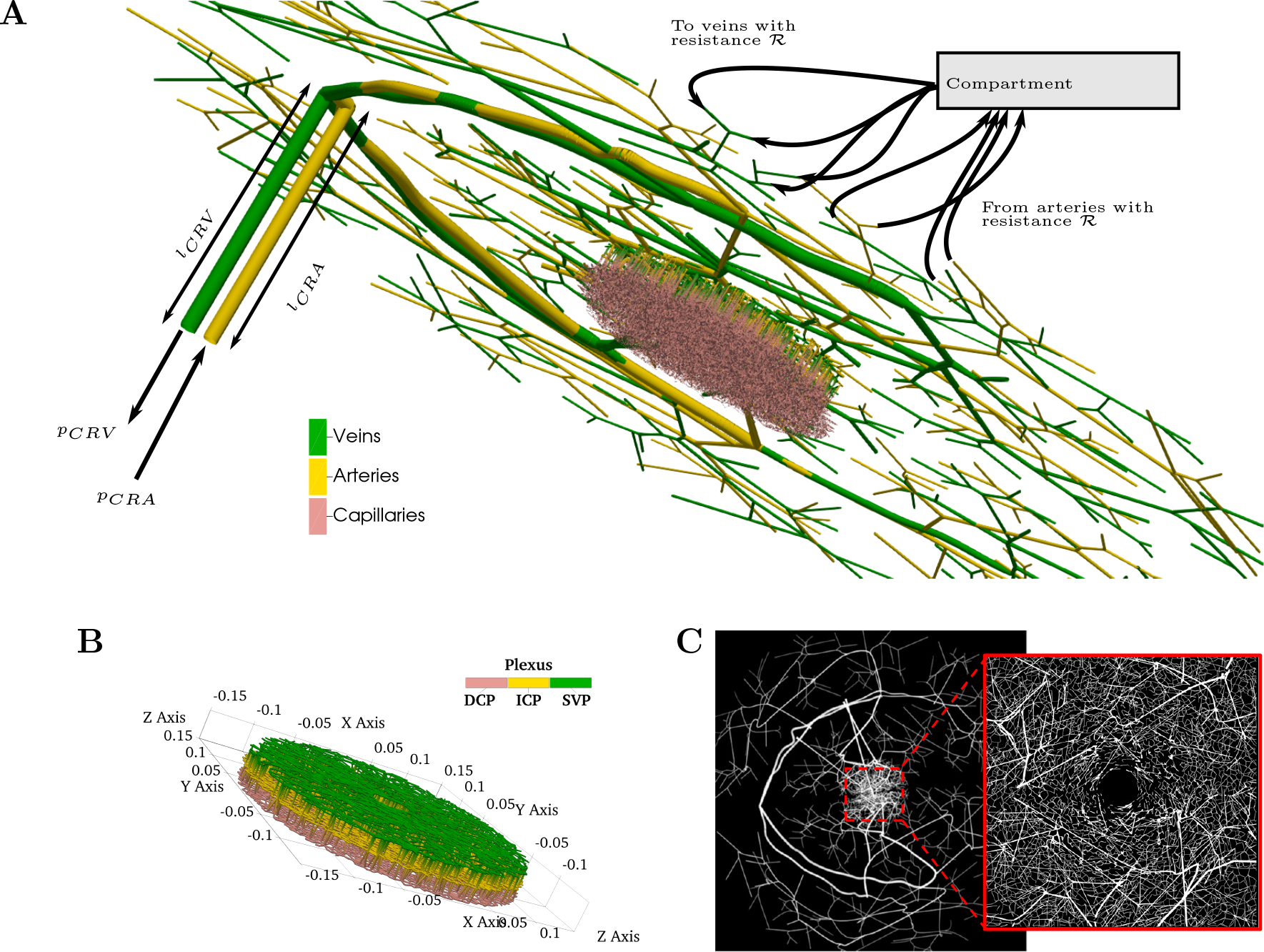
(**A**) A visualisation of one generated individual vasculature. The haemodynamics simulations use blood pressure at the CRA and CRV as inputs. Outside the macula, terminal arteries are linked to a compartment through an artificial resistance *ℛ* which redistributes the flow to terminal veins through connection with the same resistance *ℛ*. In the macula, capillaries connect arteries to veins across three vascular plexi. (**B**) Zoomed in view of the dense macular region from (**A**). (**C**) En-face view of the SVP in the temporal region and the macula (zoomed-in inset). SVP: superficial vascular plexus. ICP: intermediate capillary plexus. DCP: deep capillary plexus.

#### 2.1.1 Macrovasculature

##### 2.1.1.1 Statistical shape model

Major temporal arcade vessels were manually segmented from the DRIVE dataset^52^. The major temporal arcade vessels correspond to the four vessels (two veins, two arteries) that branch directly from the CRA and CRV and extend towards the superotemporal and inferotemporal quadrant of the retina, as shown on Figure 1A. Vessel centreline segmentations, from their branching at the level of the optic disc to the boundary of the visible retina, were extracted from 8 colour fundus photographs of eyes without retinopathy and centred at the fovea. Segmentation was performed using the ‘freehand line’ tool in Image J 1.48^50^. The pixel coordinates for each curve were extracted and translated so that the fovea is at the origin. This ensures that the model learns the distance between the optic disc and the fovea and between the arcades and the fovea, which may have importance in disease^8^. For images of right eyes, curves are reflected across the y-axis so that all shapes correspond to left eyes (i.e., the optic disc will be on the left-hand side of the image). A simple principal component analysis-based statistical shape model learns the features of all four temporal arcade vessels (see Supplementary Material S1). The generated shapes are converted to length units using the rule of thumb for fundus photographs: 10^*°*^ *≈* 5 mm, where the angle describes the field of view of the fundus camera. The generated vessels are linked to the CRV or CRA accordingly and assume the same radius as the central retinal vessels. The radius of the CRA is given in Table 1 whereas the radius of the CRV is larger by a factor 1.11^27^.

**Table 1:**
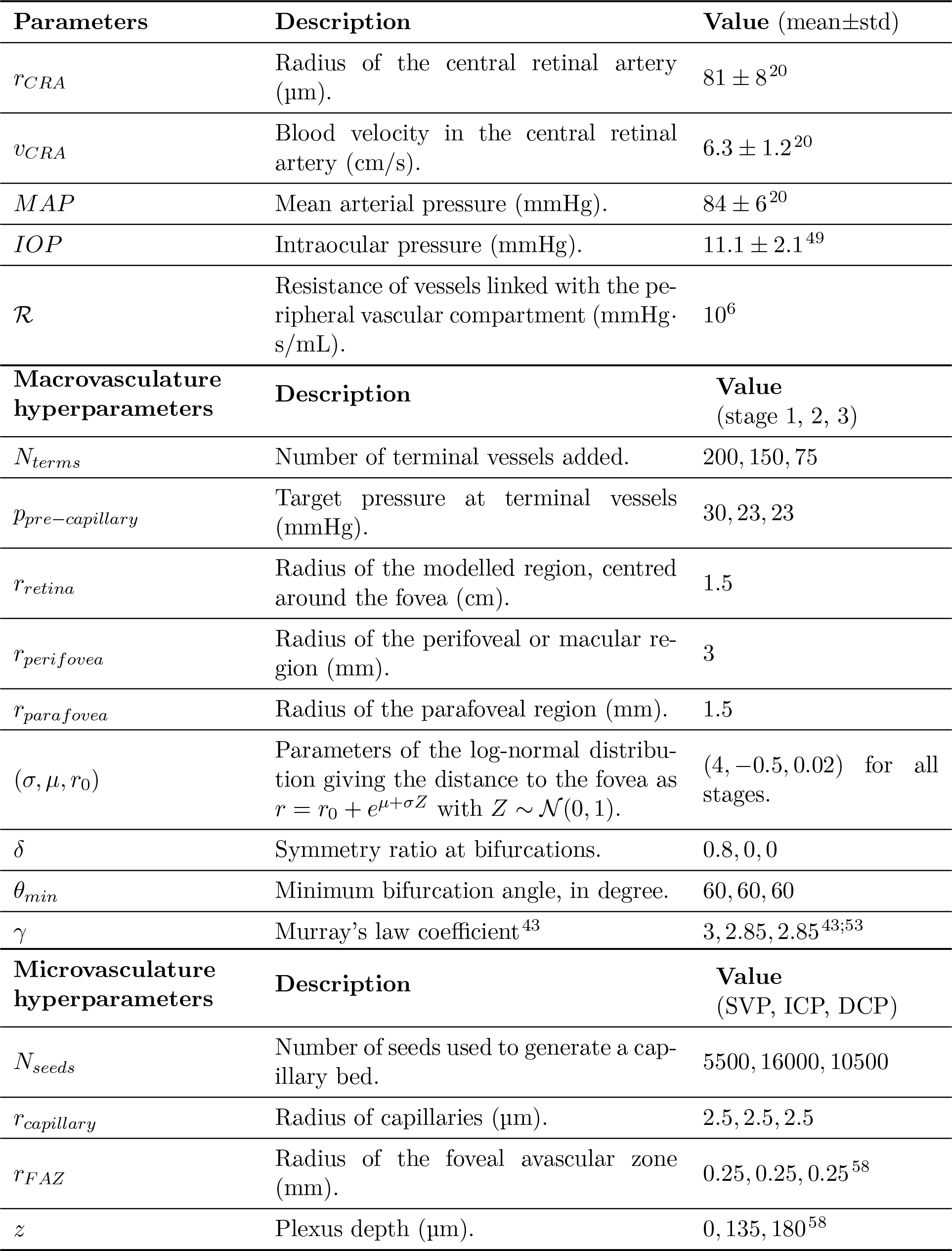
List of model parameters and their ranges.

##### 2.1.1.2 Arterial/venous branching trees

From each output of the statistical shape model, a pair of trees, one arterial and one venous, is generated.

At this stage, trees are structured branching trees using a space-filling algorithm. The algorithm is a modification of the constrained constructive optimisation (CCO) algorithm proposed by^54^. With the CCO, trees are grown to minimise the tree’s total volume while, for each addition of vessel segments, keeping a constant pressure drop from inlet to outlet and satisfying several geometrical constraints.

In addition to the original CCO’s constraints, tree growth is geometrically constrained to prevent vessels from crossing the line that passes through the optic disc centre and the centre of the fovea, which separates the superior and inferior halves of the retina. Additionally, we used a custom probability distribution to select the location of new vessel segments to mimic an angiogenic process biased towards the fovea, while keeping the foveal avascular zone (FAZ) free of vessels. Indeed, the fovea has higher concentration of cells, and therefore more metabolic needs, compared to the rest of the retina^65^. The CCO is applied in three stages, within three circular regions: in a disk of radius *r*_*retina*_, in a annulus with radii *r*_*parafovea*_ and *r*_*perifovea*_ and finally in a disk of radius *r*_*parafovea*_ (see Figure 1).

To simulate growth biased towards the fovea while keeping the FAZ free of vessels, the coordinates of a segment endpoint are given by (*x, y*) = (*r* cos *θ, r* sin *θ*), where *θ* follows a uniform distribution over the interval [0, 2*π*) and *r* follows a log-normal distribution (see Table 1). Arterio-venous networks of the superficial vascular plexus are generated in three steps: 1) the CCO is applied to create a backbone of larger arterioles and venules from the arterial and venous arcades. The arterial and venous backbones are grown separately in the first step. For each tree, the CCO requires volumetric blood flow at the root (CRA or CRV) and a pressure drop across the vasculature. Blood flow in the CRA is computed from its radius and blood flow velocity. From conservation of mass, blood flow is the same in the CRV. Ocular perfusion pressure (OPP) refers to the pressure drop between the CRA and CRV, namely:

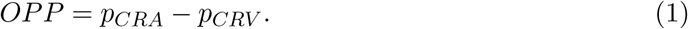

Pressure in the CRV is assumed equal to IOP^5;29;63^. Pressure in the CRA is estimated from the mean arterial pressure (MAP)^5;29;63^:

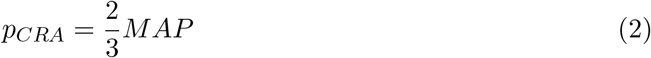

The pressure drops across the vascular trees are set to *p*_*pre−capillary*_ *− p*_*CRV*_ for the venous tree and *p*_*CRA*_ *− p*_*pre−capillary*_ for the arterial trees. The value of *p*_*pre−capillary*_ is taken from pressure in pre- and post-capillary vessels in the theoretical model by^53^.

##### 2.1.1.3 Microvasculature

At the macro scale, arterioles and venules generally follow a bifurcating structure, with parent vessels giving rise to two daughter branches. The CCO algorithm follows this logic to create vascular trees. At the micro scale, however, capillaries tend to form complex nets, forming loops and anastomoses^4^ that are incompatible with the CCO’s logic. Therefore, we adopted the methodology proposed in^35^ to generate capillary beds connecting the arterial and venous trees. In short, a disk of the size of the macula (see Table 1) is meshed with a Delaunay triangulation generated from *N*_*Seeds*_ randomly sampled points within the disk. The centroids of the triangles are used to generate a Voronoi diagram. In brief, a Voronoi diagram partitions the plane into polygonal regions centred around input points. The edges of the polygons form the capillary bed. In the SVP, capillaries coexist with arterioles and venules, but should not intersect them. Therefore, capillaries intersecting with arterioles or venules are removed from the capillary bed. This also creates a capillary-free region which is found surrounding arterioles in the SVP^4^.

In the SVP, a proportion *α* of arterioles and venules and all terminal vessels within the macula are connected to the nearest capillary. Because of the lack of specific data, *α* was arbitrarily set to 40 % in all simulations unless specified otherwise.

Inter-plexi connections remain subject to debate^4;13;16^. Since the ICP and DCP are modelled in the macula only, inter-plexi connections are based on the findings by^4^ in the parafovea. Specifically, arterioles and venules within the macular area of the SVP bifurcate to the ICP and those branches immediately bifurcate to the DCP. This corresponds to the most prevalent patterns in the histology study^4^. From the SVP, 30 % of the arterioles and venules were selected for bifurcation to the ICP and the bifurcation points were added in the middle of the selected vessels.

All capillaries are initially given the same radius *r*_*capillary*_ unless they are connected to an arteriole or venule, in which case their initial radius is twice that of other capillaries. Diameter transitions at bifurcations are smoothed using the method proposed by^35^. In short, the diameter of a segment becomes the average of the diameters of itself and of the parent and daughter branches.

The last necessary step for haemodynamics simulation is to give direction to the capillaries create from the Voronoi diagram. This can be achieved by using the diffusion equation. Representing the vasculature as a graph, values are assigned to nodes: 1 for arterial nodes, 0 for venous nodes and 0.5 for capillary nodes. The graph Laplacian of the vasculature is used to update the nodal values, creating gradients along edges (vessel segments). These gradients provide an ordering of the capillaries, from high to low value, that ensures the graph remains acyclic, which is a necessary condition for haemodynamics simulations.

### 2.2 Haemodynamics model

Blood is modelled as an incompressible, Newtonian fluid flowing in a network of connected tubes by the Hagen-Poiseuille equation. This modelling framework considers the vasculature as a arrangement of connected, straight tubes across which pressure drop Δ*p*, vascular resistance *R* and volumetric blood flow *Q* are related by:

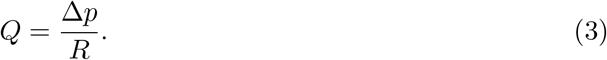

Vascular resistance is a function of the tube’s radius *r*, length *l* and the blood viscosity *μ* as follows:

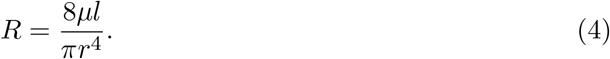

Blood is a non-Newtonian fluid and is subject to the Fåhræus–Lindqvist effect^25^ in the microcirculation. Non-Newtonian effects are accounted for by a diameter- and hematocrit-dependent, empirical, effective viscosity law^51^:

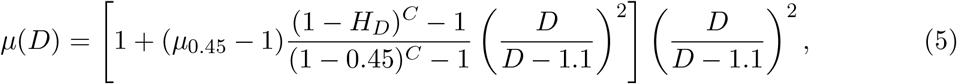

where *D* is the vessel diameter in microns, *H*_*D*_ is the discharge haematocrit,

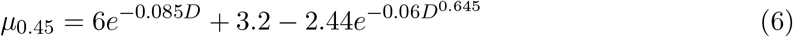

and

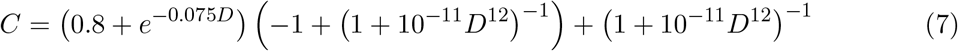

Discharge haematocrit is kept constant at 45 % in this work.

In the macular area, the vasculature is fully connected, with arterioles connecting to venules through capillaries, and therefore, no boundary conditions are required. Outside the macula, however, the first stage of the CCO detailed in Section 2.1.1.2 leaves terminal vessels with no further branches, both on the arterial and venous sides. This happens as a result of not generating the full extent of the peripheral vasculature. To close the vascular network, these terminal vessels need to somehow be linked to the CRA/CRV or haemodynamics in those vessels need to be explicitly given as boundary conditions of the model. In the absence of adequate data on haemodynamics for those vessels, we chose to instead link the terminal vessels through a compartment. This avoids the need for boundary conditions. All terminal arteries outflow into the compartment through artificial vessels with a resistance *ℛ*. All terminal veins drain the vascular compartment through artificial vessels with the same resistance. The baseline value of parameter ℛ was selected such that the haemodynamic parameters’ distribution across the vasculature is similar to experimental data^19;46^ for the baseline simulations.

### 2.3 Validation metrics

Six morphological metrics were used for comparison with OCTA:

- the four indices proposed by^17^, namely vessel area density (VAD), vessel skeleton density (VSD), vessel diameter index (VDI) and vessel complexity index (VCI);
- Fractal dimension (FD), computed with a box-counting method^39^;
- Inter-vessel distance (IVD), computed with Euclidean distance transform^36^. Only VAD and FD were used in the ICP and DCP.

For each plexus VAD was used in the model development stage to determine appropriate values for two of the hyperparameters: *N*_*terms*_ and *N*_*seeds*_ (see Table 1). Both IVD and FD require skeletonised, pixelised images of the vasculature to be computed. To create those, generated macular vessels are mapped to a white canvas, then saved as binary images.

The Euclidean distance transform was used to compute IVD and a box counting method was used to estimate FD. The remaining metrics were computed from the cumulated length (*ℒ* = Σ _*i∈𝒱*_ *l*_*i*_) and cross-section area of vessels (*𝒜* = Σ _*i∈𝒱*_ *r*_*i*_*l*_*i*_) within a given plexus and inside a field-of-view of area *χ*, as follows:

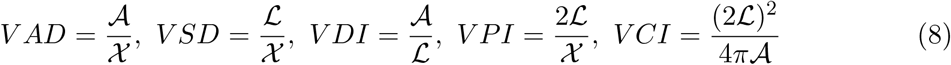

Vessels were assigned a stream order, or Horton-Strahler order. In brief, capillaries are assigned an order of 0, then moving upstream for arteries or downstream for veins, the orders are assigned as follow^4^:

- if the vessel has one branch of order *i* and all other branches are of order less than *i*, the order of the vessel is *i*;
- if the vessel has two or more branches of order *i* and *i* is the largest order among the branches, the order of the vessel is *i* + 1.

From the haemodynamics simulations, two variables were extracted to quantify macular perfusion: ‘retinal blood flow’, defined as the volumetric flow rate of blood entering the retina, and the ‘macular flow fraction’ defined as the percentage of retinal blood flow entering the macula. Spearman correlation coefficients were calculated for both haemodynamics variable and against each morphological metric. We derive 95 % confidence intervals using bootstrapping (*N* = 1000).

### 2.4 Sensitivity analysis

The method presented in this work relies on several hyperparameters listed in Table 1. The values for these parameters are either unknown (e.g., *N*_*terms*_) or are subject to uncertainty in their measurement (e.g., *δ, γ*). We performed a variance-based sensitivity analysis to decompose the variance in the model’s output (*V ar*[*Y*]). Sobol indices summarise the importance of sets of inputs with indices between 0 and 1^48^. In this work, we report first (*S*_*i*_) and total 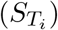 order indices, which are often enough to understand parameter importance^48^, defined as: In short, *S*_*i*_ quantifies the contribution of *X*_*i*_ alone whereas 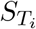 quantify its total contribution, namely its first order contribution plus all the higher-order contributions^48^. Further details can be found in the Supplementary material.

### 2.5 Uncertainty quantification

Uncertainty quantification aims at assessing the credibility of a model’s prediction^9;59^. For the haemodynamics model, uncertainty stems from two parameters, namely OPP (Equation (1)) and *ℛ*. To quantify the uncertainty brought on the correlation coefficients, the same experiment was reproduced for nine different scenarios where:

- the resistance parameter *ℛ* was set to 1 *×* 10^5^ mmHg *·* s*/*mL, 5 *×* 10^5^ mmHg *·* s*/*mL and 1 *×* 10^6^ mmHg *·* s*/*mL
- OPP was set to 80 %, 100 % and 120 % of its baseline value.

## 3 Results

A virtual population of 200 healthy vasculatures was generated. The parameters used for these simulations are listed in Table 1. The mean *±* standard deviation in OPP was 45.2*±*4.2 mmHg (range: 32.83 to 56.0 mmHg).

### 3.1 Validation of the network structure and haemodynamics

The morphology of the macula, within a disk of diameter 3 mm centred at the fovea, is compared to literature values of the same parameters computed on OCTAs^17;36;39^ in Figure 3.

**Figure 3:**
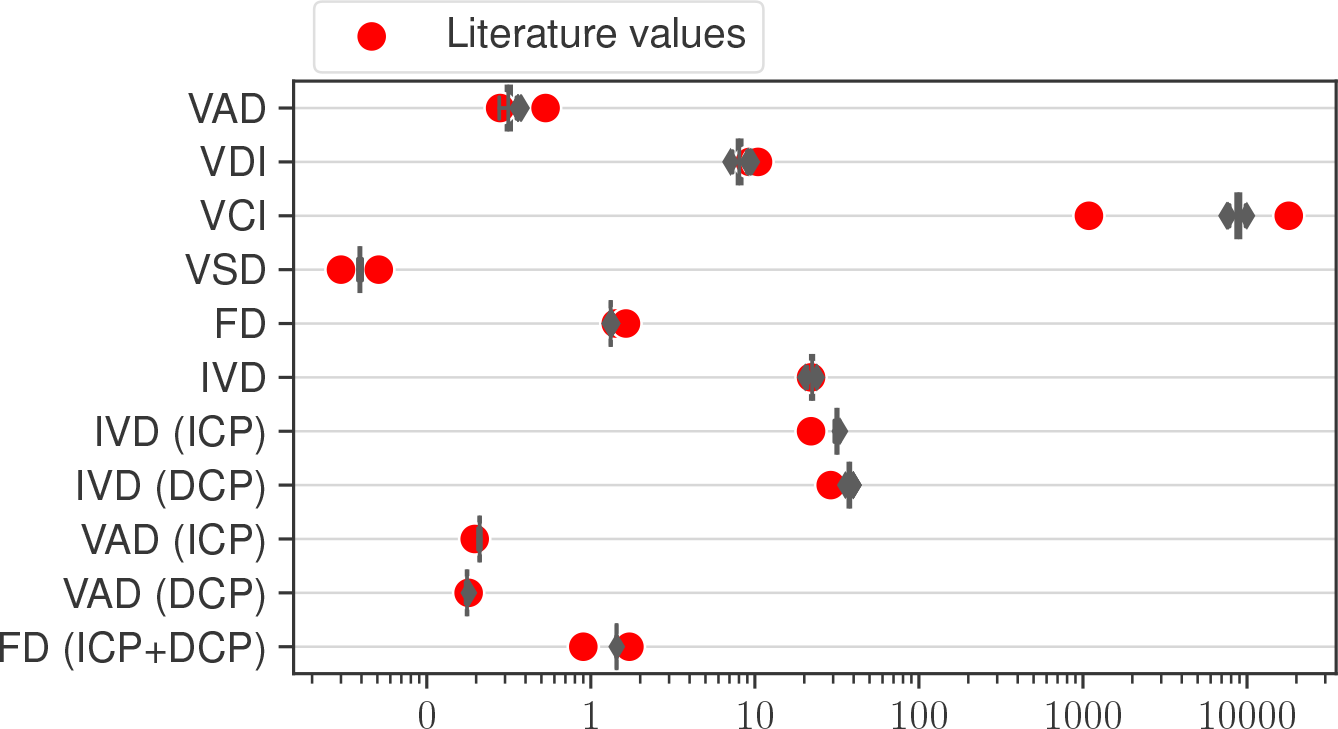
Comparison of the morphology of virtual vasculatures in the macula against OCTA measurements. Values show the deviation (in %) from the literature values (VAD^41^, VSD^41^, VDI^42^, IVD^36^, FD^1;39^ in the SVP, VAD^15^, FD^1;39^ for the ICP and DCP) for healthy eyes. Whiskers extend to the minimum and maximum within 1.5 times the inter-quartile range. Unless otherwise specified, the metrics relate to the SVP.

Figure 4 shows vessel diameters for each stream order in the macula. The distribution is similar to histological data^4^. Mean diameters were smaller in the model, but experimental data lay within the ranges of diameters in the model for each order. The ratio of average arteriole diameter to average venule diameter increased from 0.92 *±* 0.09 in order 5 vessels to 0.95 *±* 0.05 in order 1 vessels. For all orders combined, the ratio was 0.937 *±* 0.031, which is consistent with experimental measurements of 0.9 *±* 0.1^27^.

**Figure 4:**
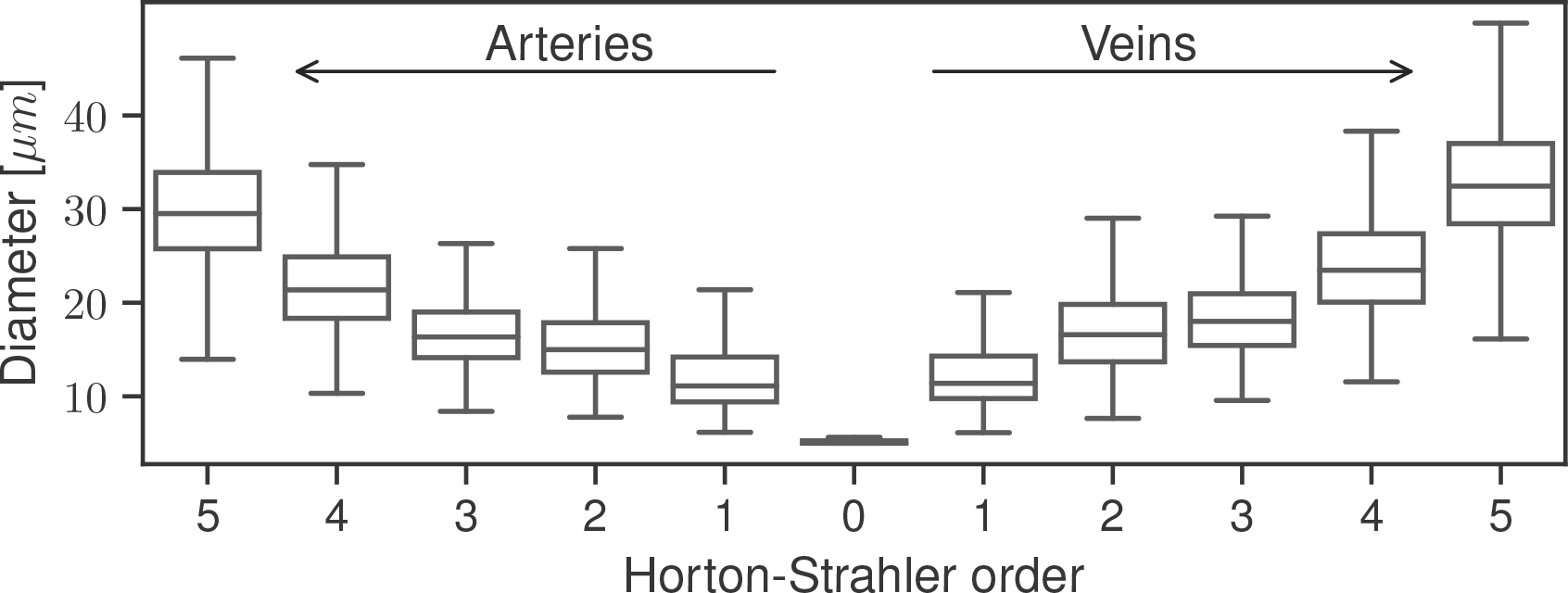
Box-and-whisker plot of the diameter of virtual vessels in the macula for each Horton-Strahler order. On average, close to 5000 vessel segments were analysed for each vasculature. Whiskers extend to the minimum and maximum within 1.5 times the inter-quartile range.

From the haemodynamics simulations, the mean *±* standard deviation for retinal blood flow, blood velocity in the CRA and macular flow fraction were 19.15 *±* 7.34 μL*/*min (range: 4.21 to 57.63 μL*/*min), 1.49 *±* 0.30 cm*/*s (range: 0.73 to 2.48 cm*/*s) and 4.52 *±* 1.19 % (ranges: 2.51 to 11.54 %). On average, retinal blood flow in the model was lower compared to experimental studies which reported a mean of 30 to 40 μL*/*min^19;46^. Blood velocity in the CRA was also lower in the model compared to the average of 6.3 cm*/*s reported by experimental work^20^. Blood velocity along the vasculature is compared with experimental studies^19;46^ in Figure 5A. Additionally, these studies reported volumetric blood flow rates against diameter. These distributions are compared with the model’s in Figure 5B. On the venous circulation, model predictions of flow and velocity were coherent with experimental data^19;46^. On the arterial side, both velocity and flow were visually lower compared to the same studies.

**Figure 5:**
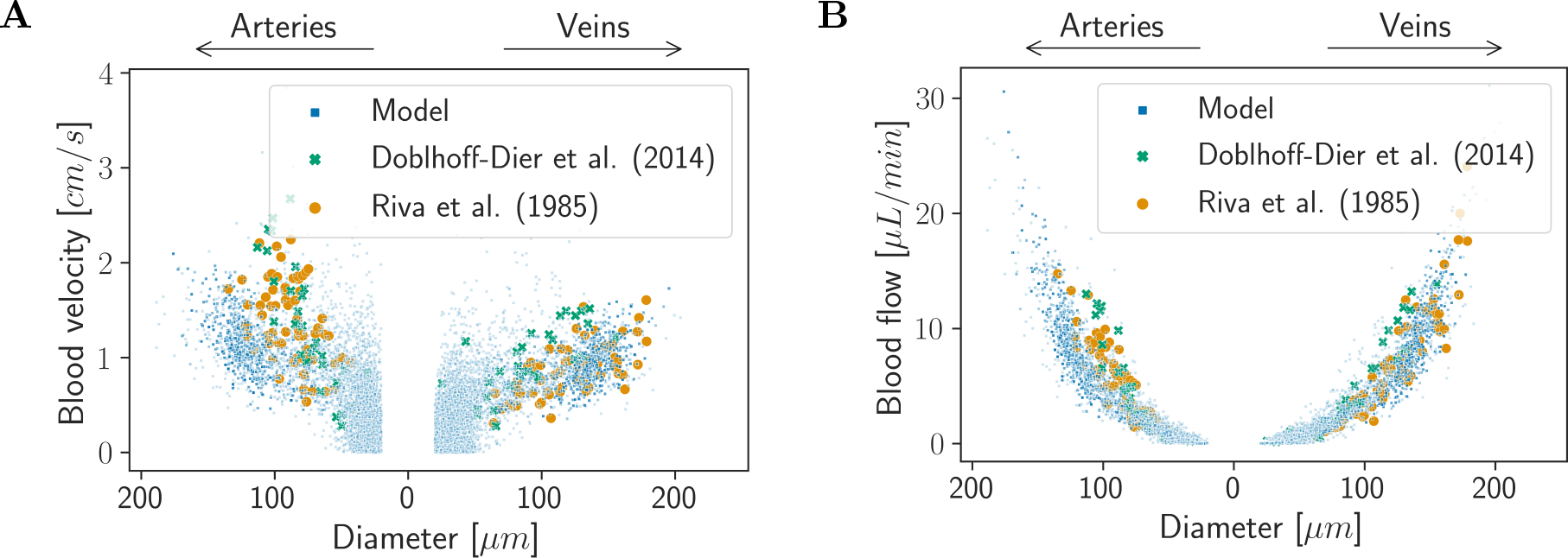
Blood velocity (**A**) and volumetric blood flow rate (**B**) distributions against vessel diameter in the virtual vasculatures compared with two independent experimental studies^19;46^.

### 3.2 Four structural variables are strongly linked to retinal function

We next look to understand how the morphology of the macular vasculature affects the haemodynamics of the retina and of the macula using our model. Figure 6 shows the Spearman correlation coefficients for each variables. Vertical lines show the threshold typically used for a correlation to be considered moderate (dotted lines) and strong (dashed lines). For the healthy virtual cohort, the model found only VAD, VDI, VCI and FD of the SVP to be predictors of retinal blood flow. The correlations with the macular flow fraction were all below 0.25, indicating weak or no correlations.

**Figure 6:**
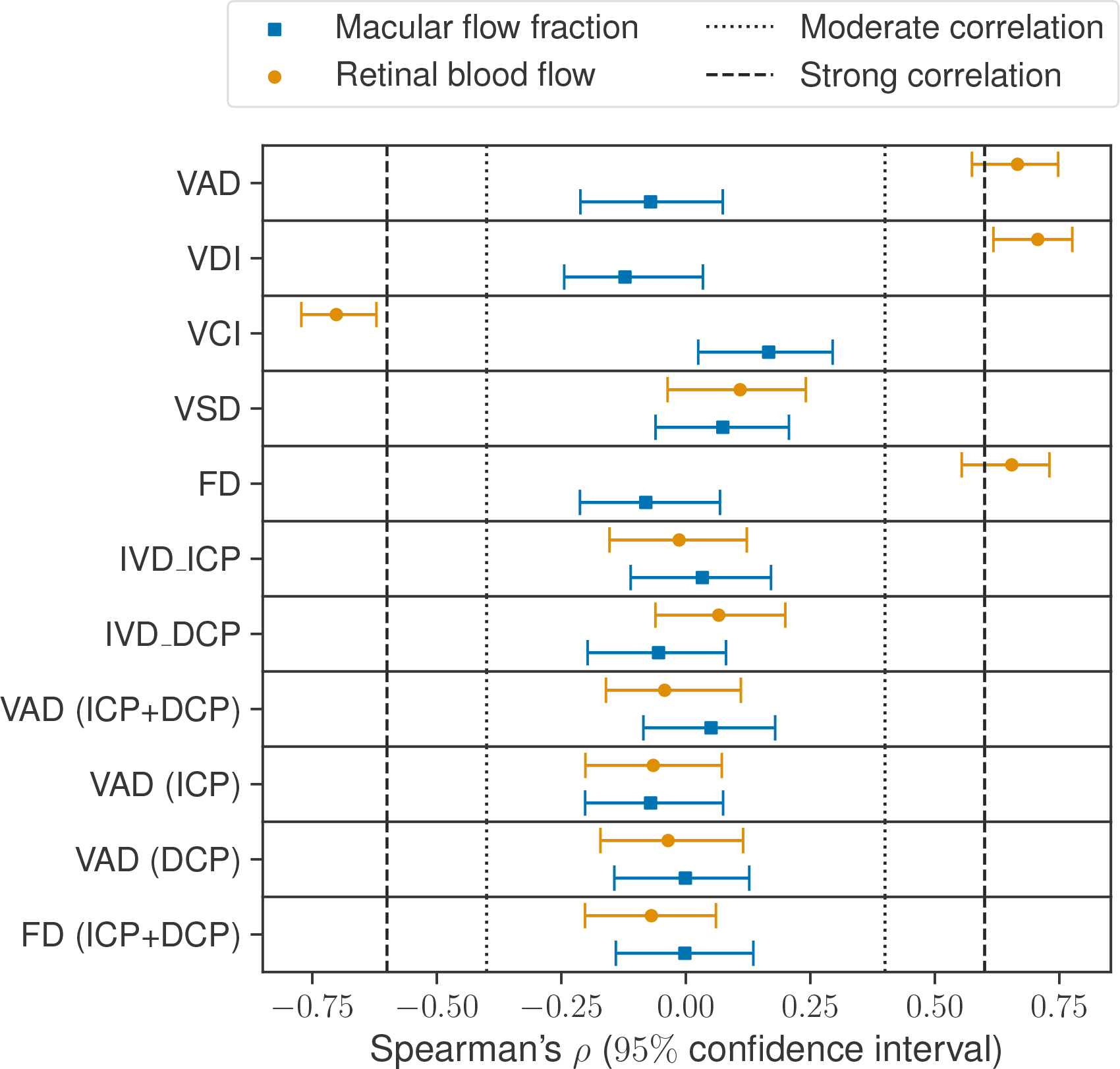
Spearman correlation coefficients testing for monotonous correlations between morphology and haemodynamics of the macula. Values closer to 1 or *−*1 indicate stronger correlations. The 95 % confidence intervals were estimated using bootstrapping.

### 3.3 Sensitivity analysis and uncertainty quantification

#### 3.3.1 Minimum branching angle dominates structural variability

Figure 7A shows the Sobol indices for 10 hyperparameters, computed with the python library SALib^31;33^ using 11 000 simulations. The parameters were sampled uniformly within the ranges presented in Table 2 using the algorithm by^47^.

**Table 2:** Ranges for the hyperparameters for the computation of Sobol indices.

| $N_{terms}$<br>stage 2 | $N_{terms}$<br>stage 3 | $l_{limfr}$ | $\delta$ | $\eta$ | $\gamma$ | $\nu$ | $\theta_{min}$ | Neighborhood<br>factor | $N_{seeds}$<br>SVP |
| --- | --- | --- | --- | --- | --- | --- | --- | --- | --- |
| [300.00, 500.00] | [200.00, 400.00] | [0.10, 0.90] | [0.10, 0.90] | [0.20, 0.50] | [2.00, 3.00] | [0.00, 72.00] | [0.00, 5.00] | [0.00, 5.00] | [800.00, 1200.00] |

**Figure 7:**
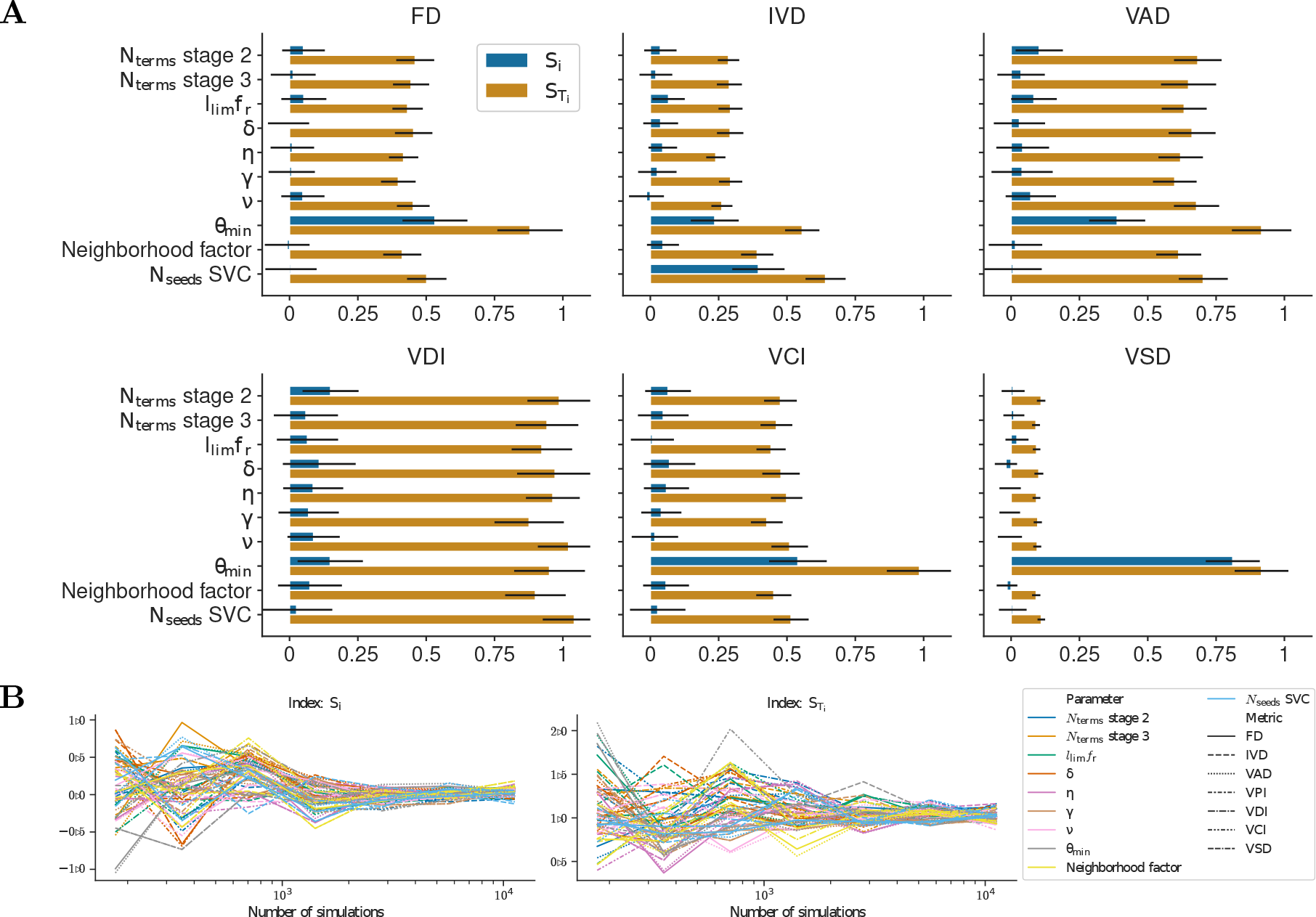
Results of the global sensitivity analysis. The bars in (**A**) show the first order (S1) and total order (ST) Sobol indices for each hyperparameters of the method with respect to each morphological metrics of the superficial vascular plexus. The error bars show the 95 % confidence interval for each indices. Convergence of the indices was checked by plotting the indices’ values with increasing number of simulations used for their computation (**B**).

For most metrics, all parameters share a similar total order and small first order. Notably, *θ*_*min*_ stands out as the most influential overall, explaining around 50 % of the variance in FD and most of the variance in 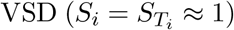 by itself. Other parameters of interest include *N*_*seeds*_ for the SVP, *N*_*terms*_ for stage 2 and, to a lesser extent, *l*_*lim*_*fr*. These results indicate that those four parameters are enough to produce a virtual population with inter-population variability, at least in the SVP.

#### 3.3.2 Macular flow fraction is independent of OPP

The results presented in Section 3 rely on several hypotheses and parameters which introduce a degree of uncertainty. Across all scenarios, the mean retinal blood flow varied between *−*26.8 % and 39.5 % of baseline values whereas macular flow fraction varied between *−*29.6 % and 22.9 %. The results for each scenario tested are shown in Table S1. Variation in OPP had almost no effects on macular flow fraction (Pearson’s *R*^2^ *<* 10^*−*3^ for all values of *ℛ*) but was linearly correlated with retinal blood flow (*R*^2^ *>* 0.98 for all values of *ℛ*). The coefficients are given for all scenarios in Table S2.

## 4 Discussion

A virtual cohort of 200 healthy patients was generated and analysed. The generated vasculature showed good agreement, structurally, with experimental and clinical measurements. The haemodynamics model showed good qualitative agreement with two independent experimental studies. The model predicted strong association between several microvascular parameters and total retinal blood flow.

### 4.1 Validation

In Section 3.1, we validated the models against experimental data. The morphology of the SVP was within the ranges of values found in the literature for healthy retinas, as quantified by OCTA^1;17;36;39;41^. In the ICP and DCP, VAD were very close to values reported in an histology study^15^ but IVD was larger in both plexuses compared to OCTA data^36^. This was more marked in the ICP compared to the DCP. The morphology of the microcirculation delineated on OCTA is sensitive to various factors, including scan post-processing^41;42^ and segmentation of the different plexuses, which makes direct comparison complicated.

The morphology of the ICP and DCP vasculature was very homogeneous across the generated cohort, as indicated by the small standard deviations. As demonstrated by the sensitivity analysis, this may be resolved by varying *N*_*seeds*_, though reasonable bounds need to be defined. Figure 4 showed that the model’s distribution of diameters across Horton-Strahler order was similar to experimental data^4^. However, capillaries were smaller in our model compared to the data. In their study,^4^ considered as capillaries any vessel with diameter *<* 8 μm, whereas in our model capillaries were assigned a diameter of 5 μm or 10 μm if they are directly connected to an arteriole or venule. This strategy may be too simplistic to represent the spread of diameters in the vascular bed. Others have suggested to update the diameter of vessels based on blood pressure from a first haemodynamics simulation^35^. More in depth analysis of the capillary beds is necessary in order to develop an appropriate strategy. In the meantime, sensitivity analysis and uncertainty quantification can help improve the reliability of the model.

Figure 5 showed that the model’s predictions of blood velocity and flow across the vasculature were coherent with experimental data. However, in the arterial circulation, both flow and velocity were slightly lower in the model^19;46^. Similarly, blood flow and velocity in the CRA were both lower in the model compared to experimental data^19;20;23;46^. As seen in Equations (3) and (4), blood flow and velocity are respectively proportional to the fourth and second power of vessel radius. Therefore, an increase in radius by a factor of 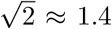 for velocity and 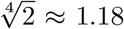 for flow would be sufficient to double the model’s predictions. It is unclear whether vessel diameter measurement in the experimental studies^19;46^ included the vessel wall in the measurement. We assumed that the diameters were those of vessel lumen, which could lead to an overestimate of lumen radii of 20 %^7^. In experimental studies, the same relation between flow and radius is assumed and blood flow is estimated from velocity *v* and diameter *D* measurements as *Q* = *vπD*^2^*/*4. Therefore, even a small measurement error in vessel diameter combined with error in measurement in velocity still results in large deviations from the true blood flow. Both measurements are challenging and prone to errors^34^. In addition, all parameters in our virtual populations were sampled from independent normal distributions which is likely an incorrect assumption as, e.g., vessel diameter is likely to be correlated with arterial pressure and IOP^21^. As discussed by^19^, studies have reported average total retinal blood flow ranging from 30 to 80 μL*/*min^19;23;46^. Despite the uncertainty in measurements, most studies seem to agree on values in healthy eyes around 30 to 40 μL*/*min^19;46^. Despite the difference in haemodynamics in the CRA, Figure 5 shows that the discrepancy with experimental data is reduced as the vessels branch out. Similar to^19^, blood velocity seem to scale linearly with diameter for larger vessels, but this trend is lost in smaller vessels. The overall lower blood velocity can be attributed, as discussed above, to the discrepancy in velocity in the CRA.

The simulations found that macular flow fraction was between 2.51 % and 11.54 %, however, to best of our knowledge, there is no data to validate these values. The macula has twice the density of cells compared to the rest of the retina^65^. Assuming that regions of higher cell density require similarly higher blood flow, it can be estimated that the macula requires 15 to 30 % of the total retinal blood flow, though these are rough estimates.

In Section 3.3.2, we quantified the effects of the two parameters of the haemodynamics model on total retinal blood flow and macular flow fraction. Increasing *ℛ* is similar to gradually closing the connections to/from the vascular compartment. Therefore, flow is shunted towards the macula and macular flow fraction increases. Decreasing *ℛ* has an opposite effect, however, the macular flow fraction will eventually reach a plateau when the CRA reaches its maximum capacity in term of blood flow: regardless of the resistance of paths outside the macula, blood will flow through the macula and total retinal blood flow is bounded by physical constraints. Indeed, in our model, for a given OPP, flow in the CRA is theoretically bounded by the radius and length of the CRA according to Equation (3). The same effect explains the non-symmetrical changes in total retinal blood flow as *ℛ* is decreased (see Figure S4).

### 4.2 Associations between structure and function

Analysis of the 200 virtual vasculatures revealed associations between several of the morphological metrics and total retinal blood flow. In particular, larger VAD, VDI, and FD in the SVP were strongly associated with larger retinal blood flow. In contrast, larger VCI in the SVP was strongly correlated with smaller retinal blood flow. Interestingly, VAD in the ICP showed a moderate positive correlation with retinal blood flow. However, VAD in the DCP and in the combined ICP-DCP complex, as well as FD in the ICP-DCP complex, did not show any significant correlations. None of the tested metrics were significantly associated with the fraction of flow transiting through the macula. Uncertainty quantification showed that those results were independent of the haemodynamics model’s parameters. As shown in Figure S3, varying OPP does not have any effects on the Spearman correlation coefficients. Increasing the value of *ℛ* shifts the value of the coefficients for macular flow fraction towards the coefficients for total retinal blood flow. When *ℛ* becomes large, macular flow becomes close to retinal flow, as seen in Figure S4, therefore increasing the impact of macular morphology on the overall system.

In Section 3.2, the correlation coefficients between morphological metrics and haemodynamics were obtained from one-to-one comparisons and therefore do not capture possible interplay between morphological metrics. Additional analysis is require to better understand which metrics or combination of metrics are strong predictors of blood flow.

### 4.3 Developing VPs with a smaller parameter space

In Section 3.3 we presented the results of a global sensitivity analysis of our method’s hyper-parameters on the morphology of the vasculature in the SVP. The results were presented as first and total order Sobol indices for 10 inputs and 6 outputs in Figure 7A. These indices were extracted from a large number of simulations in order to ensure convergence, which was reached with around 8000 simulations, as seen in Figure 7B.

The total orders indices were globally similar for all parameters and brought little explanation of the importance of parameters. Computing second order indices may be necessary to reduce the number of parameters before attempting to generate a different population or to pursue uncertainty quantification. However, second order indices require larger number of simulations to achieve convergence with Monte-Carlo methods and are therefore expensive to compute^48^. In this case, developing a surrogate model (e.g., polynomial chaos expansion) might be required^48^. Nonetheless, the first and total order indices suggests that the parameter space can be reduced to as little as four parameters, depending on which morphological metric is deemed more important.

Interestingly, *θ*_*min*_ appeared as the most influential parameter. In particular, it was the sole parameter influencing VSD. This results confirms the hypothesis by^61^ that branch geometry is correlated with VPI, which is twice VSD in Equation (8). The associations between structure and haemodynamics should be investigating further, perhaps with spatial metrics which can be compared with OCTA measurements^61^.

### 4.4 Limitations

Our method has several limitations that should be acknowledged. We did not model the peripapillary capillary plexus, ignored the curvature of the retina, and assumed a unique interplexi connection pattern. We also assumed that each plexi lay in a two-dimensional plane. Additionally, as described in Section 2.1.1.3, direct connections between arterioles/venules and capillaries in the SVP were added with a likelihood *α* which was arbitrarily set to 40 %. Similarly, the likelihood of an arteriole or venule to bifurcate to the deeper layers was also arbitrarily set to 30 %. The effects of these two values has yet to be assessed.

The effects of uncertainty in measurements (i.e., MAP, IOP, *r*_*CRA*_ and *v*_*CRA*_) on the generated vasculature remain to be quantified. The number of hyperparameters is large and global sensitivity analysis show their effects on vascular metrics to be non-local. Therefore, directly adapting the method to generate different virtual populations may prove challenging. Indeed, as stated by^2^, efficient generation of virtual populations requires knowledge of plausible ranges for the model parameters and optimising over the set of model parameters. Reducing the number of parameters to the most influential ones appear necessary and the sensitivity analysis presented in this study is a first step towards this goal.

At this stage, we have not considered the joint distribution of the population parameters such as *r*_*CRA*_ and MAP, IOP, *v*_*CRA*_. These are likely to be strongly related and ignoring these associations may create discrepancies in our model’s output when compared to experimental data. However, these joint distributions are not readily available, to the best of our knowledge, but might be inferred from different studies in the future.

The haemodynamics model proposed in this study makes several simplifying assumptions. In particular, plasma skimming effects, which lead to non-constant haematocrit, and non-Newtonian effects are important aspects of the haemodynamics in the microcirculation which were not incorporated in this model^25;51^. Additionally, we assumed that there were no lateral connections between the ICP and DCP and the circulation outside the macula. This is similar to assuming both plexi being connected ‘*in-series*’ with the SVP, which is now known to be only partially correct^4^. Finally, the parameter *ℛ* introduced in this model remains unknown and its value was based on simple computation of the estimated macular blood flow and uncertainty quantification showed that this parameter had a strong effect on the fraction of flow going to the macula. However, it had limited effect on the total retinal blood flow (at most 30 % of variation compared to baseline). In future work, its influence on other haemodynamics measurements should be thoroughly tested.

## 5 Conclusion

While macular microvascular parameters serve as potent disease biomarkers, their relationship with retinal perfusion remains ambiguous. Our method establishes a versatile framework for exploring the interplay between retinal vasculature structure and function. Designed to generate virtual populations from just four parameters and various quantitative OCTA-based measurements, the method can easily be adapted to different populations.

In our study, the model generated a population of healthy eyes, revealing robust connections between macular morphology and total retinal blood flow, independent of model parameters. The initially large hyperparameter space is effectively reduced to four hyperparameters for precise population generation.

In future work, diverse virtual populations will be created to assess model predictions in diseased maculas. This approach, complemented with haemodynamics and oxygen modelling takes an essential step towards understanding the significance of vascular imaging biomarkers and their relation to retinal diseases.

