## Supplementary material for "Linking Structure and Function: Image-Based Virtual Populations of the Retinal Vasculature" (from another version of this preprint: version 1, 2023-12-05)

### Supplementary material S1

#### Statistical shape model

The curves extracted from the fundus photographs were smooted using B-splines and their inflection points computed. For a B-spline curve $\Gamma(t) = (x(t), y(t))$ sampled at times $t_i$, $i = 1,...,k$, the inflection points are times $t_i$ where $\varphi(t) = x'(t)\,y''(t) - x''(t)y'(t)$ changes sign, i.e., $sign(\varphi(t_{i-1})) - sign(\varphi(t_i)) \neq 0$. In this work, we used $k = 30$ and $t_i = i \times k^{-1}$ to compute the inflection points. Higher sampling rates lead to more inflections points. The start and end points of the curves along with the inflection points form the primary training data-set for the statistical shape model. Because not all segmented curves yield the same amount of inflection points, we up sample the curves by splitting the longest vessel segments in half, repeating as necessary. This results in $4$ training sets, one for each arcade, with varying number of training points between sets. All shape in a training sets are aligned using the Kabsch-Umeyama algorithm [1]. Effectively, all shapes are rotated, translated and scaled as to minimise the root-mean-squared deviation between the shapes and their (aligned) mean.

For arcade vessel $j$, with $m_j$ points in each shape, a data matrix $X_j \in \mathbb{R}^{2m_j,n}$ is assembled by stacking the curves row-wise. Denoting $(e_{j,i}, \lambda_{j,i})$ the eigenvectors and associated eigenvalues of the Pearson correlation matrix of $X_j$, construct $P_j \in \mathbb{R}^{2m_j, l_j}$ by stacking the $l_j$ eigenvectors with the largest eigenvalues column-wise. The number of modes of variations $l_j$ may be chosen as to explain $95\%$ of the variance in the data. Finally, denoting $\overline{x_j}$ the mean of all shapes, one can create a new, unique shape $x$ using $x = \overline{x_j} + P_j b$ where $b$ follows a $l_j$-dimensional, centered normal distribution with standard deviation $diag(\lambda_{j,i})$, $i = 1,...,l_j$. The resulting $m_j$ points are interpolated using centripetal Catmull-Rom splines [2] before being used in the next steps.

#### Sobol indices

Let $Var[Y]$ be the variance in the model's output. For $d \in \mathbb{N}$ input parameters, $Var[Y]$ is decomposed into fractions $V_u$ attributed to sets of inputs$X_u$ as:

$$Var[Y] = \sum_{i=1}^{d} V_i + \sum_{1 \le i < j \le d} V_{i,j} + \dots + V_{1,2,\dots,d},$$

where

$$V_i = Var[E(Y|X_i)],$$

is the first order contribution of variable $X_i$ to $Var[Y]$ and

$$V_{i,j} = Var[E(Y|X_i)] - V_i - V_j,$$

Is the contribution of the interactions between $X_i$ and $X_j$. For higher interactions (e.g., $V_{i,j,k}$), similar expressions can be built.

In this work, we report first $S_i$ and total $S_{T_i}$ order indices defined as:

$$S_i = \frac{V_i}{Var[Y]},$$

$$S_{T_i} = \frac{E[Var(Y|X_{\tilde{i}})}{Var[Y]},$$

where $X_{\tilde{i}}$ is the set of all input parameters excluding $X_i$ and $V_i$ is the first order contribution of variable $X_i$ to $Var[Y]$ [3].

### Uncertainty quantification

**Table S1 - Results from the 9 scenarios tested for uncertainty quantification.**

| | | **Total retinal blood flow ($\mu L/min$)** | | **Macular flow fraction (%)** | |
|---|---|---|---|---|---|
| $\mathcal{R}$ | OPP | mean | std | mean | std |
| $10^5$ | 80 | 17.9 | 7.3 | 3.2 | 1.0 |
| | 100 | 22.4 | 9.1 | 3.2 | 1.0 |
| | 120 | 26.9 | 11.0 | 3.2 | 1.0 |
| $5 \times 10^5$ | 80 | 15.3 | 5.9 | 4.5 | 1.2 |
| | 100 | 19.1 | 7.3 | 4.5 | 1.2 |
| | 120 | 23.0 | 8.8 | 4.5 | 1.2 |
| $10^6$ | 80 | 14.0 | 5.2 | 5.5 | 1.3 |
| | 100 | 17.5 | 6.5 | 5.5 | 1.3 |
| | 120 | 20.9 | 7.8 | 5.5 | 1.3 |

**Table S2 - Pearson's $R^2$ scores testing linear association between $\mathcal{R}$ and OPP for the 9 scenarios tested for uncertainty quantification.**

| **Dependent variable** | $\mathcal{R}$ | | | **OPP** | | |
|---|---|---|---|---|---|---|
| **Group** | $OPP = 80$ | $OPP = 100$ | $OPP = 120$ | $R = 10^5$ | $R = 5 \times 10^5$ | $R = 10^6$ |
| **Total retinal blood flow** | 0.82 | 0.82 | 0.82 | 0.89 | 1.00 | 0.99 |
| **Macular flow fraction** | 0.93 | 0.93 | 0.93 | 0.0 | 0.00 | 0.00 |

.

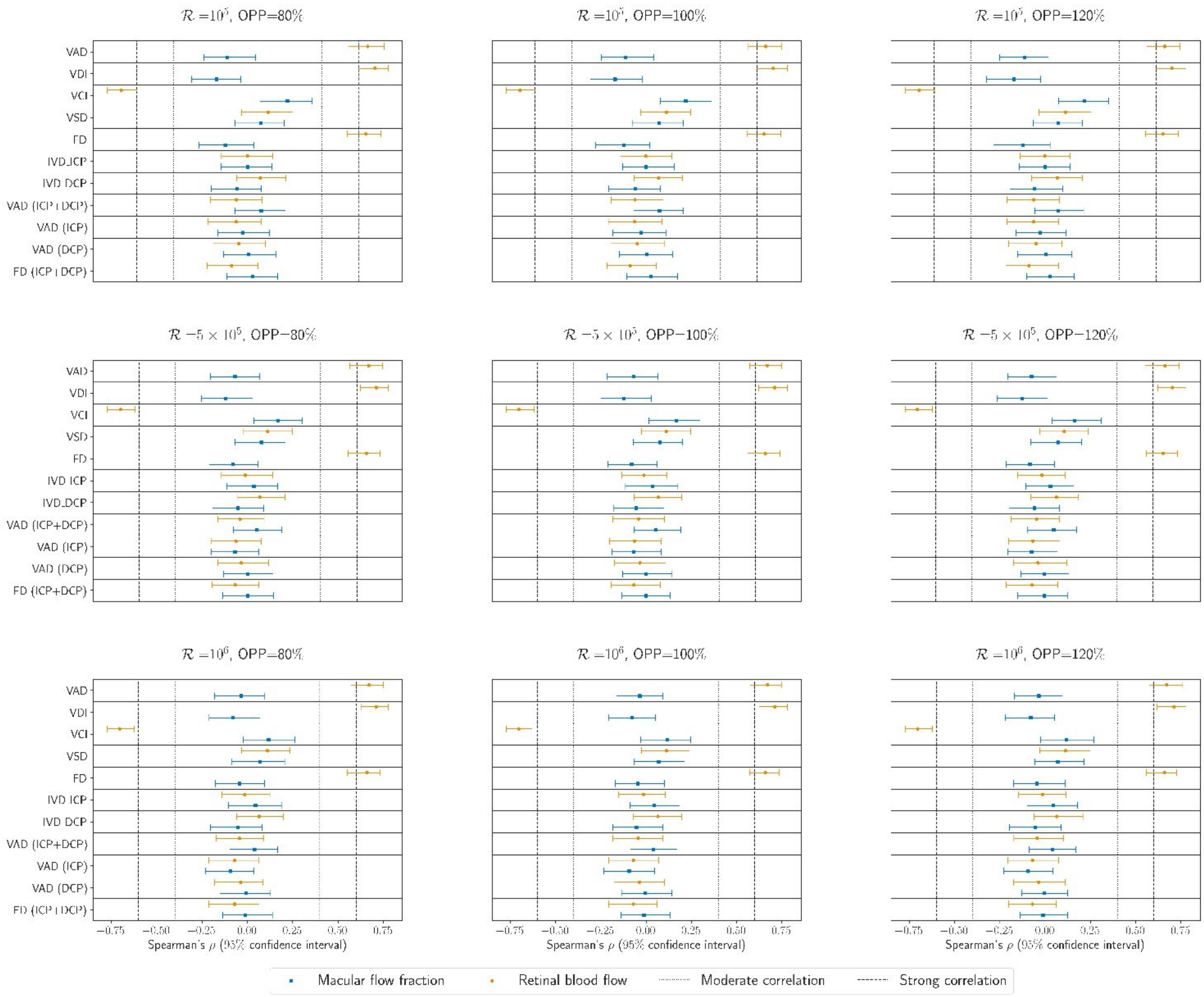


**Figure S1 - Effect of varying OPP and $\mathscr{R}$ on the associations between structural and haemodynamics outcomes.**

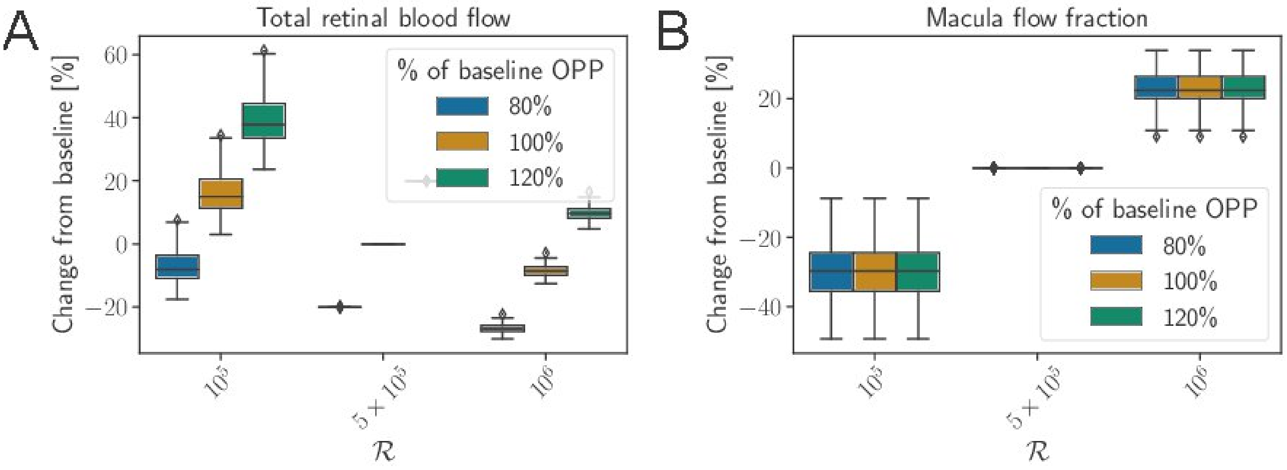


**Figure S2 - Effects of varying OPP and $\mathscr{R}$ on the haemodynamics outcomes.**
